# In vitro efficacy of Fasting Mimicking Conditions combined with targeting of starvation escape pathways against prostate cancer cells

**DOI:** 10.1101/2025.08.27.672470

**Authors:** Mahshid Shelechi, Maura Fanti, Giacomo Guiliani, Valter Longo

## Abstract

Prostate cancer (PCa) remains a leading cause of cancer-related death among men, particularly due to the development of treatment-resistant disease such as castration-resistant prostate cancer (CRPC)^1^. Emerging evidence suggests that metabolic interventions like the fasting-mimicking diet (FMD), which promotes differential fasting responses in normal and cancer cells, can enhance the efficacy of conventional therapies and overcome resistance mechanisms. We investigated the effects of FMD in vitro on androgen-sensitive and androgen-insensitive PCa cell lines, and evaluated its potential to synergize with hormone therapies and pathway-specific inhibitors. Fasting mimicking medium (FMM), a medium which substitutes the FMD *in vitro*, significantly reduced cell viability across multiple PCa models, and sensitized them to agents targeting PI3K-AKT-mTOR signaling and cholesterol biosynthesis, including rapamycin, simvastatin, pictilisib, and alpelisib. RNA sequencing of C4-2 cells under FMM conditions revealed upregulation of cholesterol biosynthesis genes and key escape pathways, identifying novel vulnerabilities that could be exploited therapeutically. Notably, combined inhibition of androgen signaling, PI3K pathways, and cholesterol synthesis in FMM conditions resulted in potent cytotoxicity, while also requiring lower doses of each agent. Our findings underscore the therapeutic potential of integrating FMD cycles with molecular-targeted therapies in PCa. Future studies should explore personalized, biomarker-driven approaches leveraging transcriptomic data to predict and exploit FMD-induced escape pathways.

## Introduction

Prostate cancer survival and proliferation are mostly driven by androgen binding to androgen receptors (AR) and activating pro-tumor pathways^2^. Even in the absence or low levels of androgen, androgen signaling remains active as evident by high expression levels of androgen-related genes such as PSA in androgen-insensitive tumors^2^. Androgen deprivation therapy (ADT) for localized disease, and ADT combined with chemotherapy for metastatic disease, can increase the overall survival by a few years, however ultimately PCa progresses to castration-resistance despite low levels of androgen^3^. Currently there is no cure for castration-resistant prostate cancer (CRPC). The crosstalk between AR and growth factor signaling pathways is well established as a driver for disease progression: growth factor receptors and their downstream kinases such as PI3K-AKT are up-regulated during different stages of PCa, and they can either phosphorylate androgen receptors directly or sensitize them to the low levels of androgen or promote growth via androgen-independent pathways^4,5^. In addition, insulin and IGF levels are associated with an increased PCa risk since they induce cell proliferation and promote androgen-independent growth^6^. Leptin, another factor that can induce cell proliferation, enhances AKT phosphorylation and increases Cyclin D1 in LNCaP PCa cells via up-regulation of the PI3K signaling pathway^7^. Fasting mimicking diets (FMDs) which in mice provide approximately 10-50% of normal calorie intake for 4-5 days, induce differential stress resistance which protects normal cells against cytotoxic chemotherapy by increasing their resilience^8^ while sensitizing to therapy cancer cells, which fail to adapt due in part to oncoprotein activity^9^. The protective effect of fasting/FMDs against the cytotoxicity of chemotherapeutic agents is partly regulated through the reduction of glucose and/or circulating IGF-1 and oncogene signaling^8,10,11^. FMD also decrease the growth and promotes the death of different cancer types by reducing many of the factors implicated in PCa including insulin, leptin, ferritin, etc. For example, we showed that the FMD decreases estrogen receptor transcriptional activity and enhances estrogen receptor inhibition in hormone receptor positive breast cancer^12^. In this HR^+^ breast cancer model, FMD cycles increase the anti-cancer effect of endocrine therapy (fulvestrant targeting ER signaling and Palbociclib targeting CDK4/6), delays the resistance to these agents and cause tumor regression. Importantly, the FMD also reverses the acquired resistance of breast cancer cells against Palbociclib and Fulvestrant ^13^.

In a recent study, RNAseq analysis of triple negative breast tumors in mice undergoing cyclic FMD identified starvation escape pathways pointed to a novel therapeutic approach: blocking these escape pathways (PI3K, AKT, mTOR, etc) combined with the FMD lead to tumor regression and reduced toxicity of inhibitors to mice^14^. In fact, the chronic use of the inhibitors cocktail without the FMD caused 100% lethality to the mice.

Clinical trials have shown that combining pharmacological PI3K-AKT and/or mTOR inhibitors and AR blockage treatments increase overall survival of prostate cancer patients^15^ but, as in mice, induce hyperglycemia causing poor tolerability. Current prostate cancer therapies are associated with a number of side effects ranging from hyperglycemia, osteoporosis, increased risk of heart disease, and fractures to Parkinson’s disease^16^. These side effects reduce the patient’s quality of life, and lead to complex therapeutic management requirements. Instead, in mice and possibly in patients, FMD prevents the increased in glucose/insulin in response to PI3K-AKT and mTOR inhibition and consequently avoids hyperglycemia^14^. In addition, FMD cycles have been shown to reduce a number of other treatment-related side effects^13,14,17–21^. FMD has been shown to modulate cellular damage and tumor microenvironment to promote T cell-dependent killing of breast and melanoma cancer cells^18,22–24^. Here, we aim to test a unique therapeutic approach that combines the FMD with hormone therapies to target the AR and other oncogenic signaling pathways in prostate cancer. We performed RNA sequencing analysis to identify the most common escape pathway responses to develop a novel escape pathway inhibitor cocktail that compliments the FMD with the goal of slowing the initiation and progression of prostate cancer but also preventing and reversing drug resistance.

## Result

### Effects of in vitro fasting on the viability of androgen-dependent and androgen-independent PCa cell lines

We studied whether androgen-dependent and independent PCa cells cultured under fasting conditions, referred to as fasting mimicking medium (FMM, 0.5g/L glucose, 1% fetal bovine serum [FBS]), will display decreased viability. Our results indicate that in vitro fasting/FMD reduces cell viability of both androgen-dependent (LNCaP) and androgen-independent (C4-2 and 22RV-1) PCa cell lines (**Fig. 1**). Notably, many of our studies indicate that this *in vitro* method is an excellent model predicting *in vivo* responses.

**Fig. 1.**
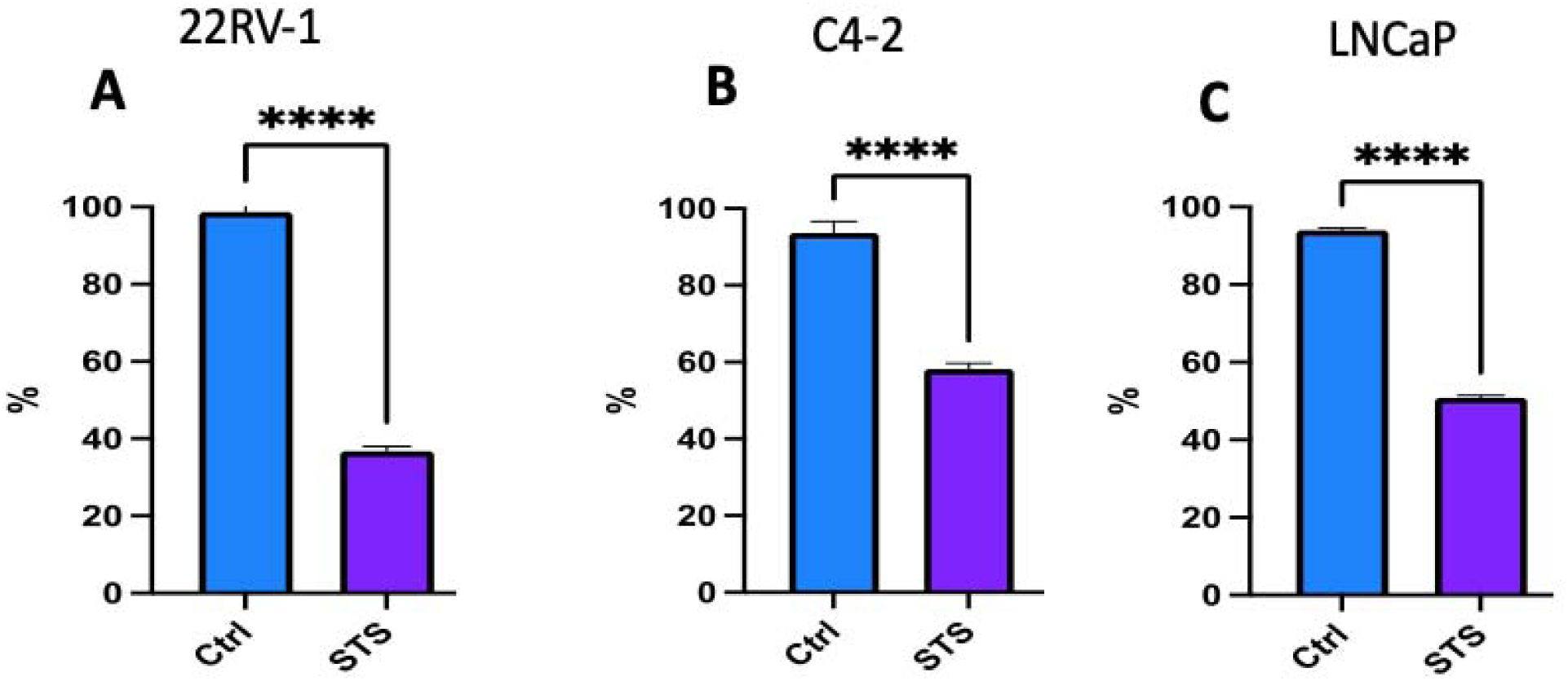
*In vitro* fasting reduces PCa viability. 72 hours of fasting (1% FBS, 0.5g/L glucose) reduces viability of **A**) 22RV-1, **B**) C4-2, and **C**) LNCap PCa cells. **** p< 0.001.

Both LNCaP and LAPC4 cell lines are androgen-dependent, and PSA producing xenografts. LNCaP has a single point mutation on a ligand binding domain of androgen receptor gene, but LAPC4 expresses a wild type AR. Castration resistant C4-2 cell lines are androgen-independent sublines of parental LNCaP cells that have been generated by several passages in vitro and in vivo ^25^. LNCaP and C4-2 cell lines were purchased from ATCC and LAPC4 cell line was gifted from Dr. Henning at UCLA. 22RV-1 is a castration resistant prostate cancer cell lines and it was gifted for Dr. Cohen at USC.

To determine if glucose reduction drives the effect of fasting mimicking medium on cell viability, cells treated with or without FMM condition with different concentration of glucose and serum. Four different cultures include: HGHS: 10% FBS+ 1 g/l glucose, LGLS: 1% FBS + 0.5 g/l glucose, LGHS:1% FBS + 1.0 g/l glucose, HGLS: 10% FBS + 0.5 g/l glucose. After 24, 48, 72 and 96 hours, cells viability was measured with both MTS and Crystal violet. Our results indicate that both androgen sensitive, LNCap, and androgen insensitive, C4-2 and 22RV-1, depend on glucose more than serum for proliferation, as adding back serum to the medium did not affect the cell viability (**Fig. 2**). Each experiment was done in triplicate.

**Figure 2.**
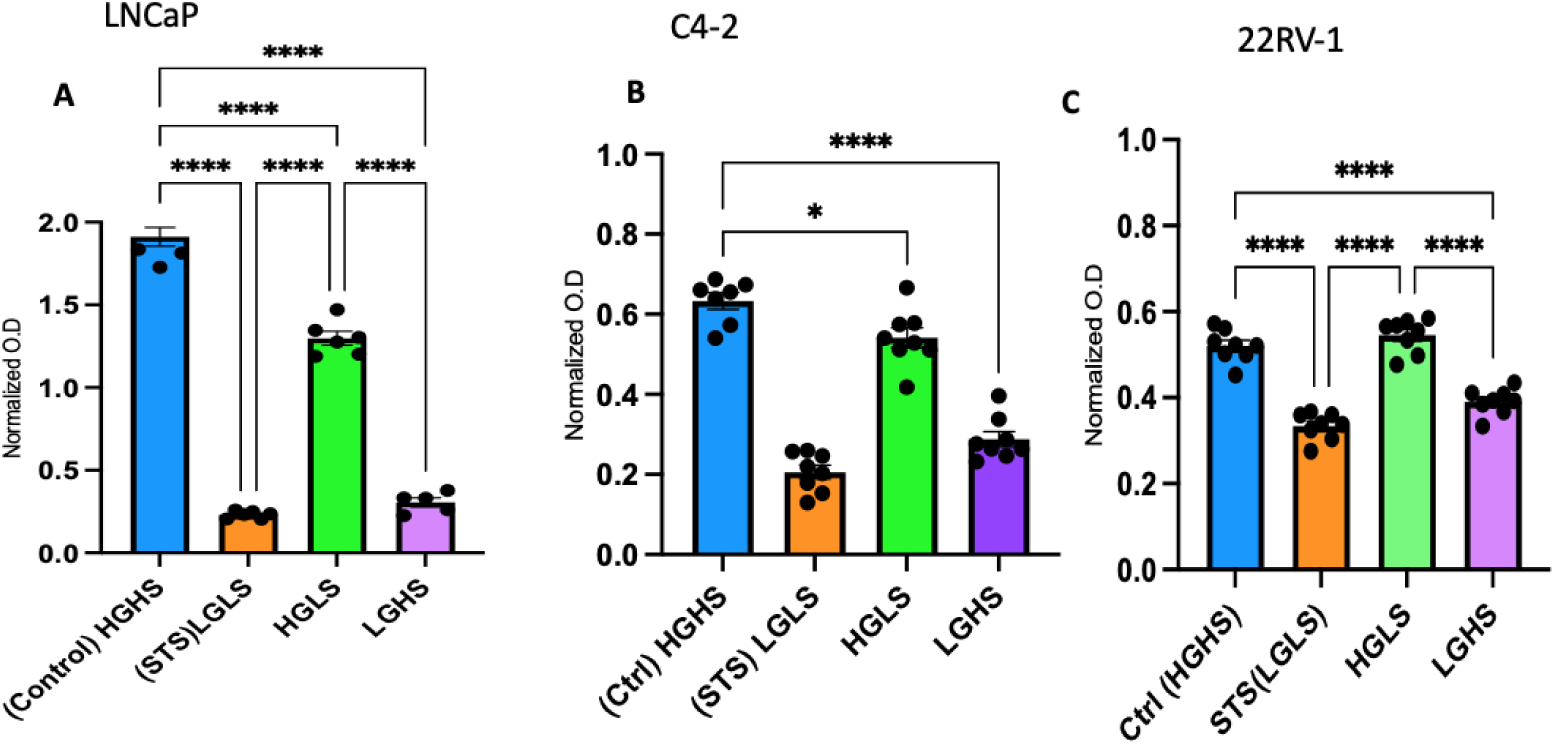
In Vitro fasting reduces cell viability via reduction in glucose rather than serum. Prostate cancer cells were seeded in 96 well plates and cultured for 72 hours with or without STS conditions, low glucose low serum (LGLS), low glucose high serum (LGHS) and control (HGHS) conditions. Thereafter, cell viability was determined. **A**) LNCaP , **B**) C4-2, and **C**) 22RV-1 PCa cells. **** p< 0.001.

### FMD activates starvation escape mechanisms that can be targeted low toxicity or non-toxic drugs

The pharmacological inhibition of the PI3K-AKT and mTOR pathways potentiates the effect of the FMD: *in vitro*, combining fasting with pictilisib, ipatasertib, and rapamycin (selective inhibitors for PI3K, AKT, and mTOR, respectively) results in enhanced cancer cell death and reduction of mammosphere numbers in SUM159 cells, partly due to a synergistic effect on the PKA signaling pathway^14^. Prostate cancer studies showed that simultaneously targeting PI3K-AKT and AR by combining respectively rapamycin and Enzalutamide results in the reduction of PCa growth^26,27^. Therefore, we tested if fasting conditions could sensitize the prostate cancer cells to the mTOR inhibitor, Rapamycin, known for its anti-aging properties^14,28,29^. Prostate cancer cell lines, C4-2 and 22RV-1, were seeded in 96-well plates and cultured with or without fasting conditions, 4 nM Rapamycin (this concentration was chosen based on the dose response experiment) or their combination for 48 hours and then cell viability and proliferation was detected (**Fig. 3**). Each experiment performed in triplicate. Our finding suggests that fasting mimicking medium (FMM) might synergizes with rapamycin to inhibit cell viability and decrease in cell proliferation in prostate cancer cell lines.

**Figure. 3.**
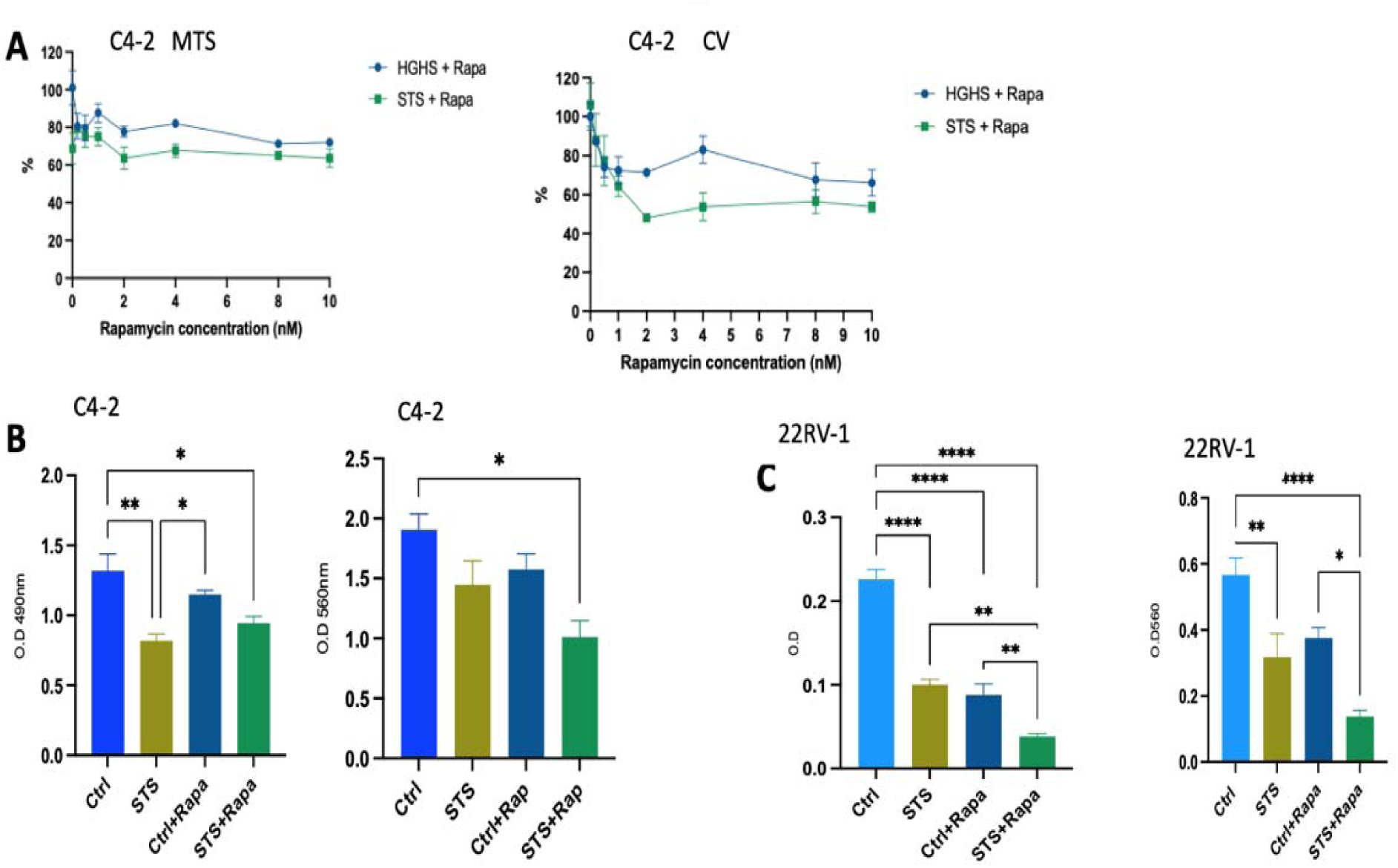
A) Fasting mimicking medium/STS, sensitizes C4-2 and 22RV-1 cells to Rapamycin. C4-2 cells were treated with Rapamycin (range 0-10 nM) for 72 hours. Cell viability was determined by MTS and crystal violet (CV) assay. **B)** C4-2 cells were grown in CTRL (1g/l glucose, 10% FBS) or fasting STS (0.5 g/l glucose,1% FBS) media. At 24h cells were treated with Rapamycin (4 nM). Cell viability was assessed by MTS and crystal violet (left to right). **C)** 22RV-1 cell viability with or without fasting STS media treated with 4nM rapamycin for 48 hours. MTS (left), crystal violet (right).

The activation of cellular escape pathways can allow differentiated cancer cells to acquire resistance to various treatments modalities, even under fasting/FMD conditions^13^. Thus, the identification of escape pathways that can be targeted to increase treatment efficacy can have a key role in cancer cell toxicity. In mouse models, Salvadori et al. performed RNA-seq analysis in SUM159 breast tumor masses from control or FMD-fed animals to identify escape pathway targets. RNA-seq analysis indicates that FMD cycles upregulate PI3K-AKT and mTOR pathways and down-regulate CCNB-CDK1 while upregulating CCND-CDK4/6 signaling axes. As anticipated, the pharmacological inhibition of the PI3K-AKT and mTOR pathways potentiates the effect of the FMD: *in vitro*, combining fasting with pictilisib, ipatasertib, and rapamycin (selective inhibitors for PI3K, AKT, and mTOR, respectively) results in enhanced cancer cell death and reduction of mammosphere numbers in SUM159 cells, partly due to a synergistic effect on the PKA signaling pathway^14^.

Identifying escape pathway mechanisms that are differentially expressed based upon fed or fasted conditions provides a unique opportunity to apply a variety of drugs, incl. those that are traditionally outside the scope of oncology. This approach of drug repurposing typically requires reduced time and lower risk over *de novo* drug discovery processes since approved drugs have already undergone all relevant phases of development for their originally intended indication^30^. Therefore, we predict that we can utilize escape pathway targets identified by RNAseq to repurpose existing drugs to further optimize PCa treatment efficacy. I performed bulked RNAseq analysis to identify the escape pathway in androgen-independent prostate cancer cell lines, that cause therapy resistance due to metabolic plasticity. C4-2 cell lines were seeded in 96 well plates. After 24 hours, cell were grown in CTRL (1 g/l glucose, 10% FBS) or fasting/FMM (0.5 gr/l glucose, 1% FBS) media. After 24 hours, cell were treated with rapamycin 4uM. The day after, total RNA was isolated using the miRNeasy Mini Kit. Libraries for RNAseq were prepared following the manufacturer protocols for transcriptome sequencing with the Illumina NextSeq 550DX sequencer (ILLUMINA). Our RNA-seq analysis indicates an up-regulation of several biosynthetic pathways with the most notable being sterol and cholesterol biosynthesis and metabolism, macroautophagy, as well as the PI3K-AKT, mTOR signaling cascade. With Volcano plot I showed that genes related to sterols/cholesterol/steroid biosynthesis pathways such as; TM7SF2, INSIG1, LSS, HMGCS1, FDFT1, MSMO1, HSD17B7, IDI1, SREBF2, LDLR, SQLE, MVK and SULT2B1 were upregulated with fasting vs. control (**Fig. 4**).

**Fig 4.**
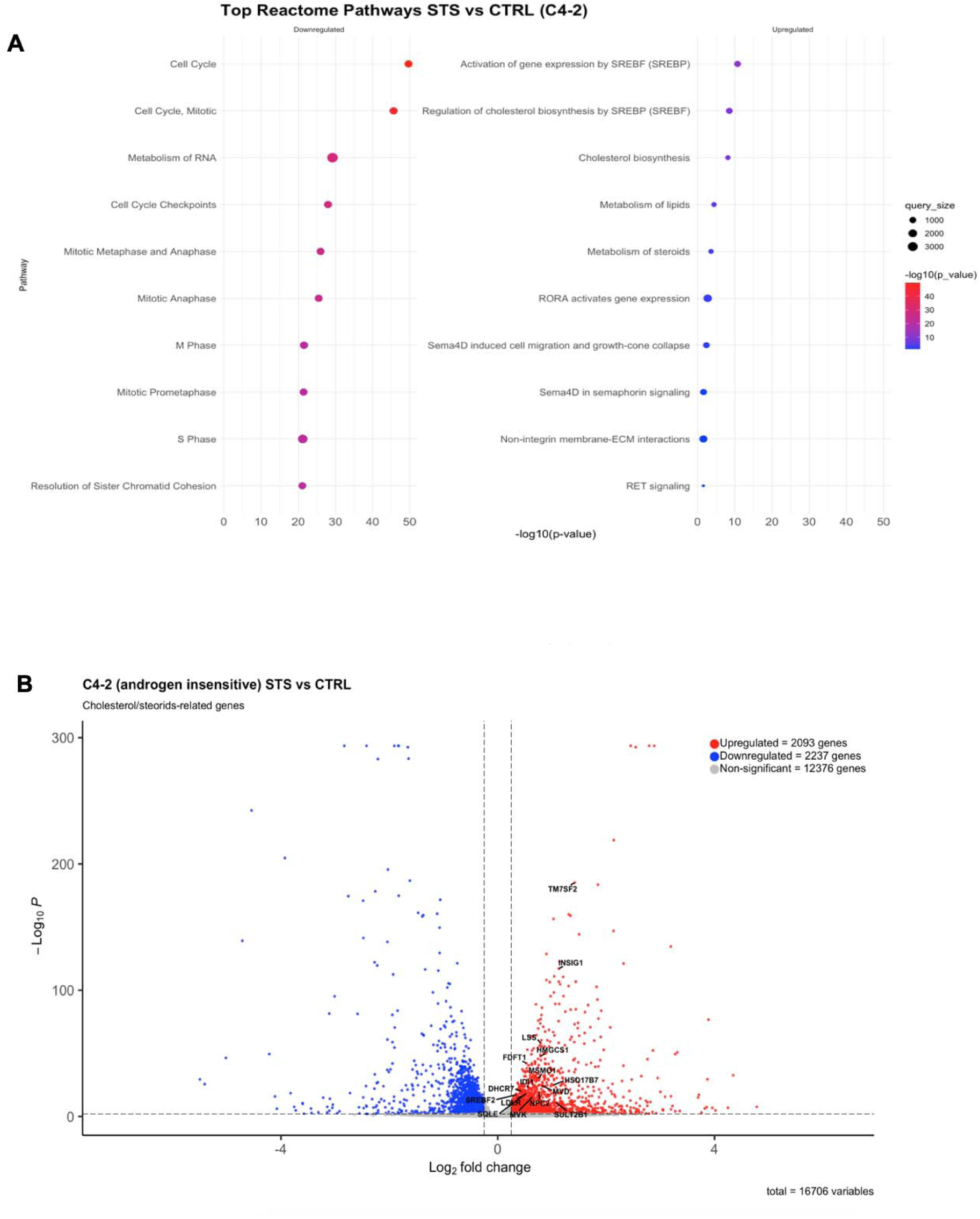
RNAseq analysis to identify escape pathway mechanism. **A)** Gene set enrichment results indicating top Reactome pathways in fasted vs. control C4-2 cells. **B)** Volcano plot of the differential gene expression in C4-2 PCa grown in control or STS media. Highlighted are the components of cholesterol/steroid genes.

In fact, it was recently demonstrated that fasting-like cell culture conditions and cholesterol biosynthesis inhibitors (CBIs) cooperate to induce anti-tumor effects in a broad range of tumor types; independent of the limited cholesterol availability due to reduced FBS concentrations in the fasting-like media^31^. Statins, cholesterol synthesis inhibitors, are widely used in cholesterol lowering therapy. In recent years, statin’s anticancer utilities have been investigated in different types of cancer such as breast cancer, multiple myeloma, lung cancer, metastatic colorectal cancer and leukemia^32^. Simvastatin, another very low toxicity drug, is a 3-hydroxy-3-methylglutaryl-CoA reductase (HMGCR) inhibitor, is widely utilized and its one of the most prescribed statin in the management of cholesterol level and cardiovascular diseases. Therefore, we focused on studying the effects of simvastatin on prostate cancer cell function in vitro. First, we performed drug tests to determine the half-maximal inhibitory concentration (IC50) of simvastatin for several prostate cancer cell lines ( C4-2, LNCap and LAPC-4) ranges from 0-120 uM with or without STS (**Fig. 5**). The in vitro concentrations of simvastatin used in studies investigating its effect on prostate cancer cells ranges from 1uM to 100uM. Our results indicated that hormone insensitive prostate cancer cell line, C4-2, is more sensitive to simvastatin (IC50=20uM) when exposed to fasting condition compared to the hormone-sensitive cell lines, LNCap and LAPC-4 which have an IC50 ∼45uM. Prostate cancer is a heterogeneous disease, therefore, simvastatin may have different effects when its combined with fasting mimicking medium (FMM) in the early vs. progression stage of prostate cancer.

**Fig 5.**
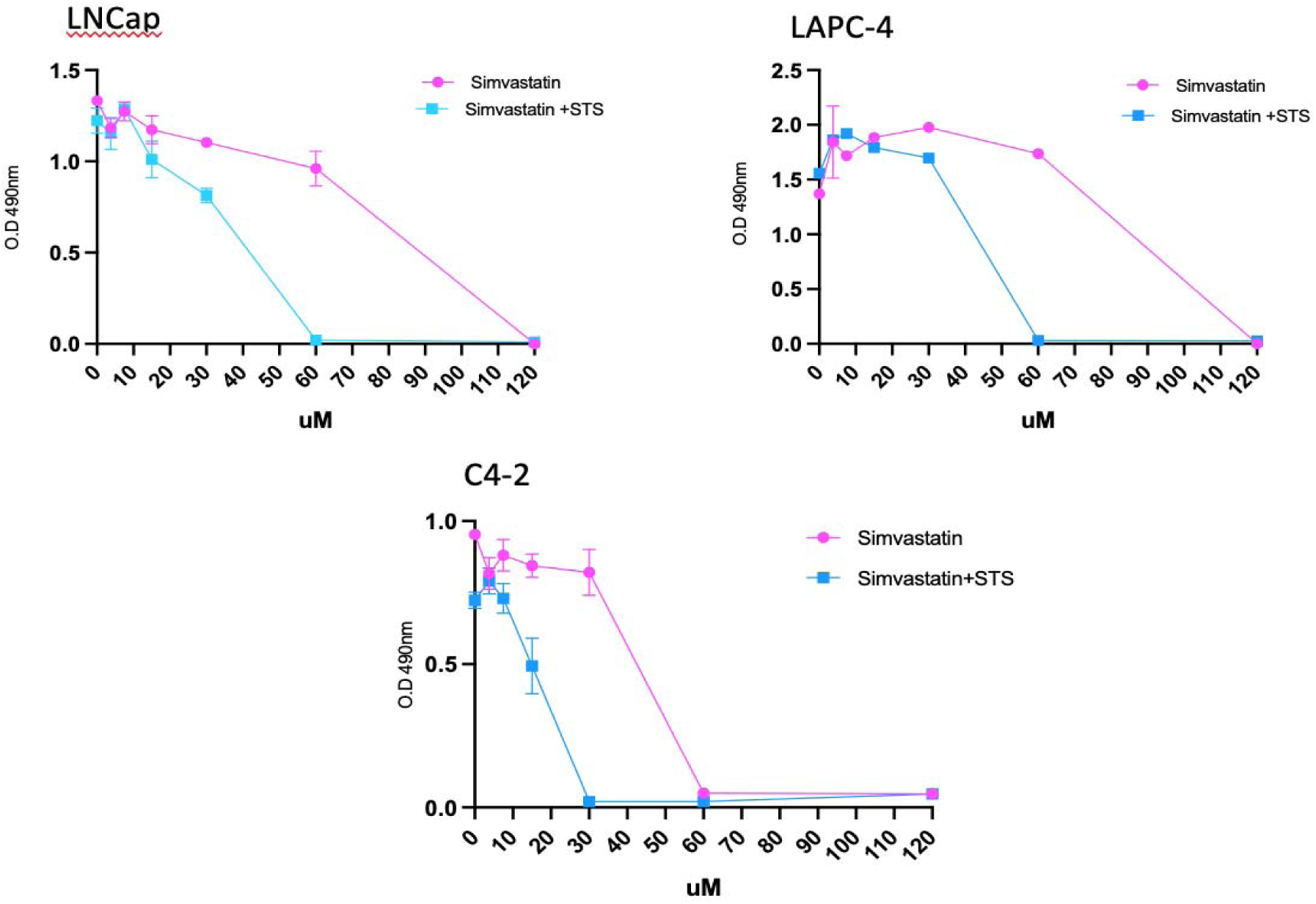
Hormone sensitive Pca cell line is more sensitive to simvastatin when combined with STS. LNCap, LAPC-4 and C4-2 prostate cancer cell lines were seeded in 96-well plates and after 24h exposed to a gradient of drug concentrations of simvastatin for 48 hours, ranging from 0-120uM. Cell viability wa**s** assessed by MTS.

**Fig.6** shows almost 0% viability when C4-2 cell lines are incubated in fasting medium and treated with only 20uM of simvastatin. In agreement with this finding, we showed that fasting mimicking conditions confer a remarkable synergistic cytotoxic effect in combination with simvastatin, in both hormone-sensitive and hormone-insensitive prostate cancer cell lines as shown a reduction of cell viability to almost 0% in both LNCap and C4-2 Pca cell lines (**Fig.7**).

**Fig. 6.**
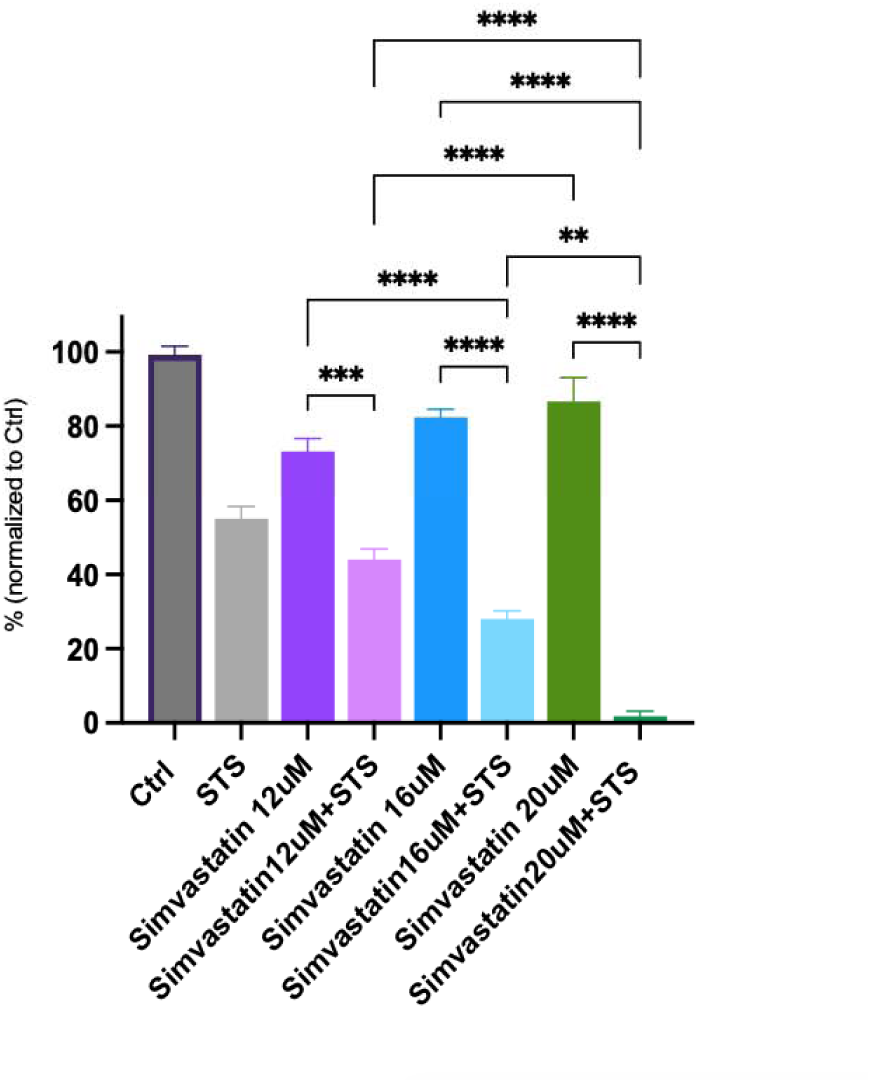
Fasting medium-induced non-oncological drug sensitization of prostate cancer cells. C4-2 prostate cancer cell viability w/ or w/o STS and treated with vehicle, or the cholesterol biosynthesis inhibitors, simvastatin, at different concentrations, 12uM, 16uM and 2OuM during the last 48h of incubation. Ceils viability was assessed with MTS.

**Fig 7.**
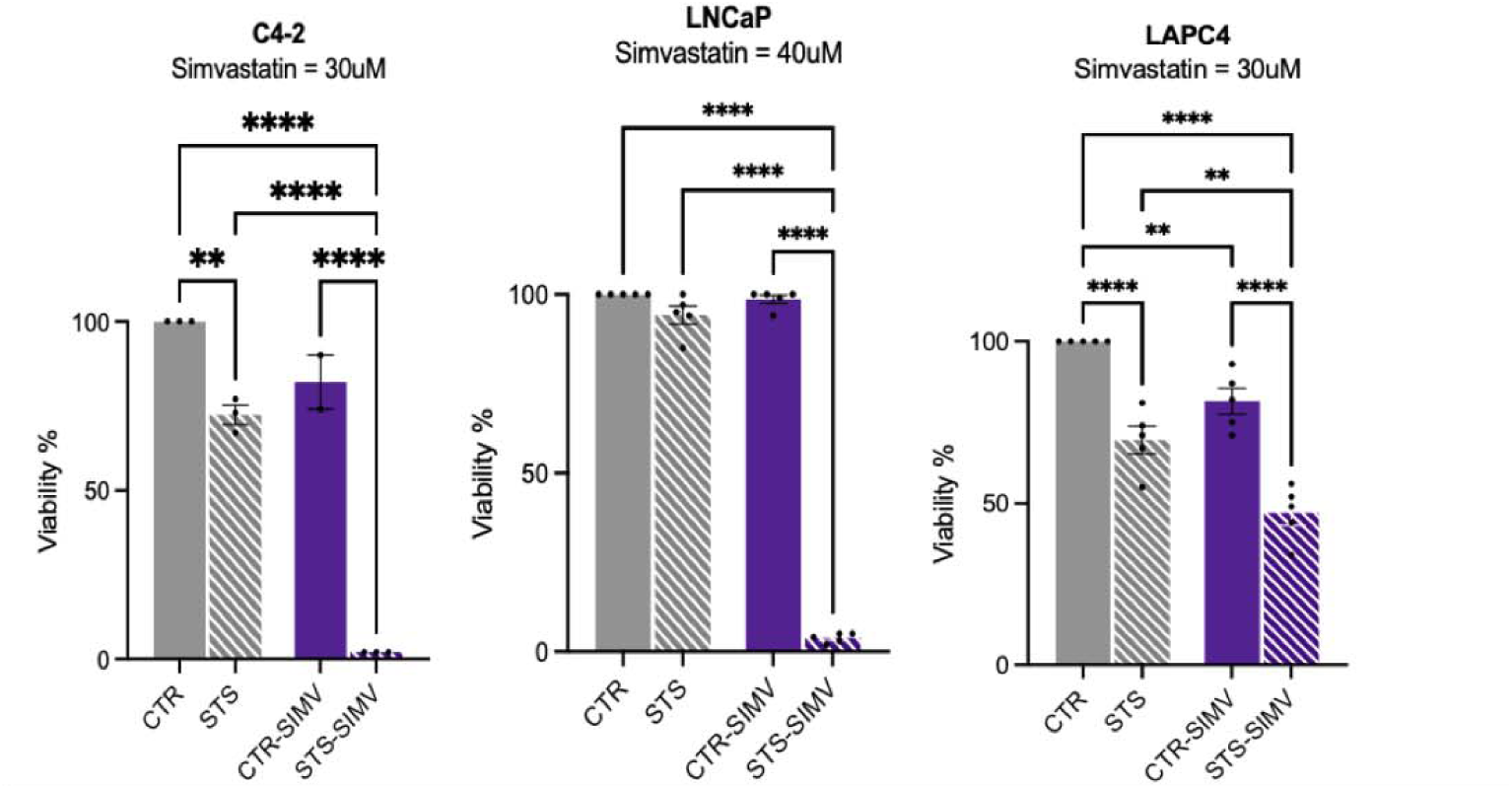
HMGCR inhibitor cooperate with fasting mimicking medium/STS to kill prostate cancer cells. Pea cells, C4-2, LNCap and LAPC-4 were treated for 72h w/ or w/o STS and for the last 48h w/ or w/o simvastatin 3OuM, 4OuM and 3OuM respectively. Thereafter, cell viability was measured by MTS. ”*‘ p< 0.001. ’

### Identify and target escape pathways in combination with in vitro fasting

Next, we tested the hypothesis that combining drugs that target these biosynthetic pathways as well as standard-of-care drugs will cause a major increase in treatment efficacy, as seen with the improved therapeutic outcomes reported for SUM159 breast cancer cells^13^. Some in vitro and in vivo studies have shown the additive or synergistic inhibitory effect of simvastatin in prostate cancer cells combined with Enzalutamide^33^. Even some human studies have shown an additive effect of statins combined with Enzalutamide or Abiraterone^34^. Here, we investigated if the selective targeting of the escape pathways identified in our RNAseq in combination with *in vitro* fasting and/or therapeutic interventions Enzalutamide can maximize treatment efficacy. One androgen-sensitive and two androgen-insensitive cell lines were plated in 96-well plates at 20,000 cells/well, respectively, in complete media in triplicates. The next day, cells were treated in fasting-like and normal growth conditions for a total of 48 hours. At 24h cells were treated with Simvastatin, HMG-CoA reductase inhibitor and Enzalutamide, Androgen receptor inhibitor, for 24 hours, as single, double, or triple treatments (*concentrations based on preliminary experiments*). Cell viability was assessed by MTS solution. In this experiment, we showed the combination of HMGCR inhibitor simvastatin and anti-androgen drug Enzalutamide has a synergistic effect on the proliferation of C4-2, LNCaP and LAPC-4 Pca cells (**Fig 8**).

**Figure 8.**
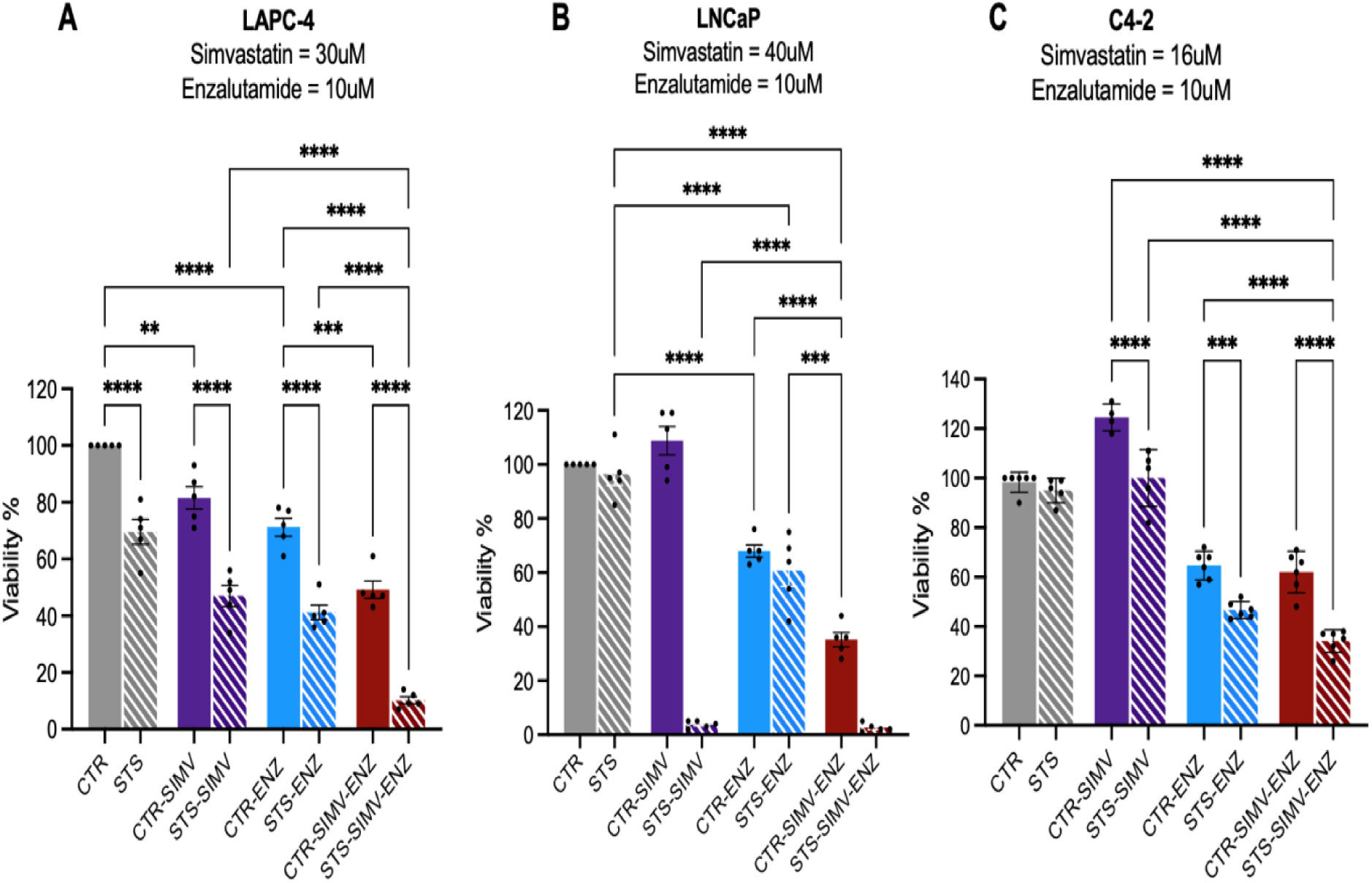
Fasting mimicking medium/STS sensitizes the prostate cancer cell lines to. **A)** LAPC-4, **B)** LNCap, and C) C4-2 to Enzalutamide when combined with the non-oncological drug Simvastatin.

To confirm the effect of fasting condition/STS on AR signaling, we analyzed expression of AR proteins. Surprisingly, the combination of STS and simvastatin decreased AR protein expression, as compared with fasting mimicking medium alone, although there was no statistical significance (**Fig 9**). A study by Kochuparambil et al. showed a relationship between Akt inhibition and inhibition of tumor growth by simvastatin (in vivo and in vitro)^35^. In fact, our results show that protein expression of phosphorylated AKT decreased when the cell lines were treated with simvastatin (**Fig 9**). Therefore, simvastatin might decrease AR protein expression when its combined with fasting mimicking medium through inhibition of AKT pathway or may reduce pAKT by down-regulating AR/AR signaling. In line with reduction of AKT phosphorylation and mTOR activity, treatment with simvastatin reduced p70S6K. Cell lines treated with PI3K inhibitor, Pictilisib, showed reduced phosphorylation of AKT and of p70S6K (an mTOR target).

**Fig. 9.**
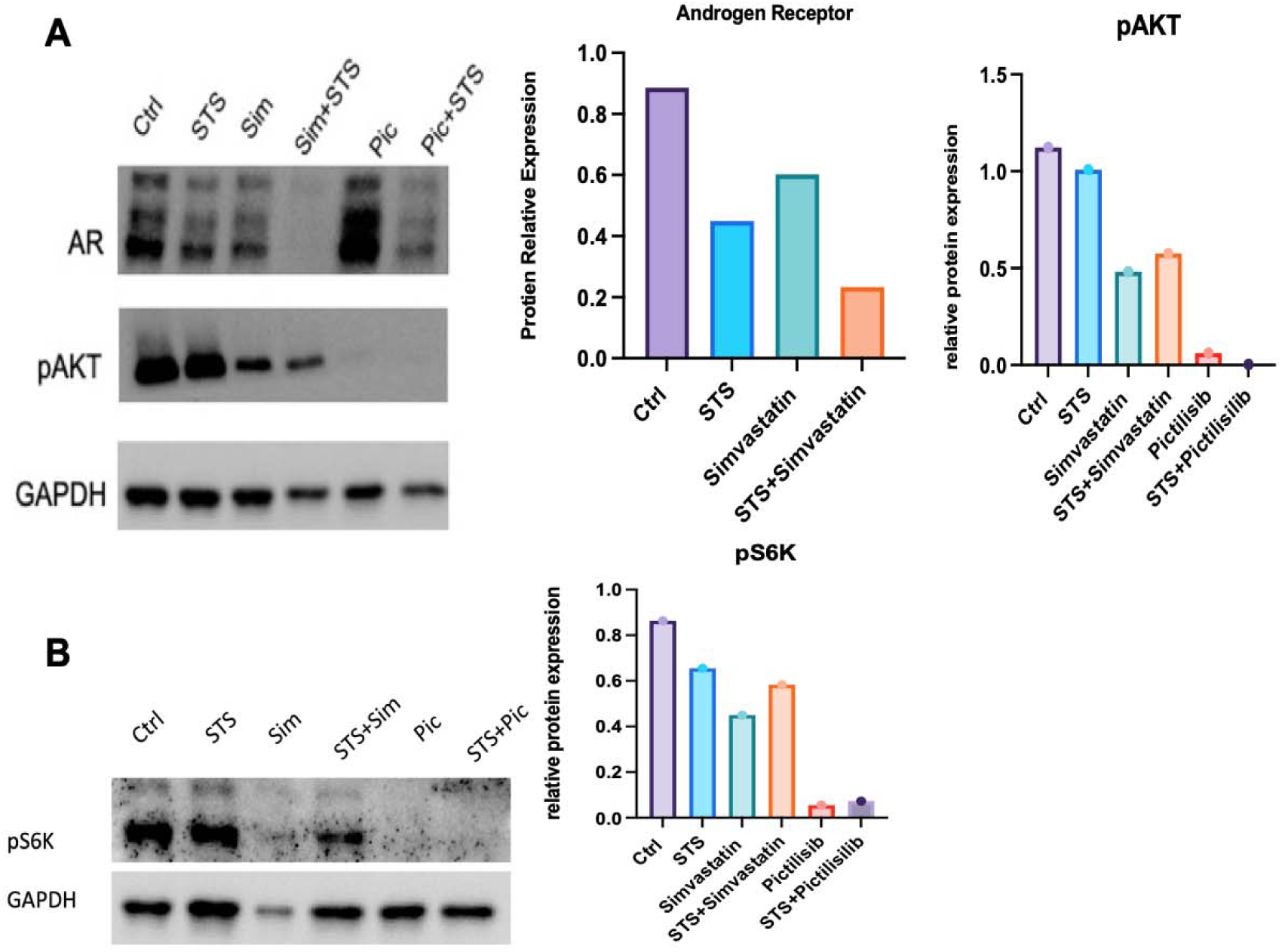
Fasting mimicking medium/STS reduces AR expression in C4-2 Pea and decrease PI3K-AKt-mTOR signaling when its combined with Pictilisib. A,B, C, C4-2 cells were seeded in 6-well plates and cultured for 48h w/ or w/o STS conditions in the presence or absence of Simvastatin 3OulVI and Pictilisib 2OuM. Thereafter, protein subjected to protein lysate generation, AR, pAKT, pS6K, as well as GAPDH, levels were detected by WB.

RNAseq analysis further indicated that in androgen independent C4-2 Pca cells, grown in fasting mimicking media, there is a upregulation of PI3K signaling pathways (**Fig.10**). We investigate if that the PI3K-AKT, mTOR signalling, and cholesterol biosynthetic pathways are required for survival by several prostate cancer cell lines in response to fasting conditions. PI3K-Akt is a key pathway activated in CRPC associated with poor prognosis^36^. Targeting of Akt with AZD5363 as monotherapy has pro-apoptotic and anti-proliferative activities in enzalutamide-resistant cell lines *in vitro* and decreases tumor growth in Enzalutamide-resistant xenografts^37^. The combination of AZD5363 and Enzalutamide decreases cell proliferation and induces cell-cycle arrest and apoptosis in PCa cell lines LNCaP and C4-2. Notably, the combination of AZD5363 plus Enzalutamide, but not either therapy alone, results in the regression of castrate-resistant LNCaP xenografts without recurrence ^37^. In a phase I clinical trial in patients with mCRPC, Enzalutamide combined with AZD5363 was considered safe and tolerable and provided anti-tumor activity with all responding patients harboring aberrations in the PI3K-AKT-mTOR pathway^38^.

**Fig. 10.**
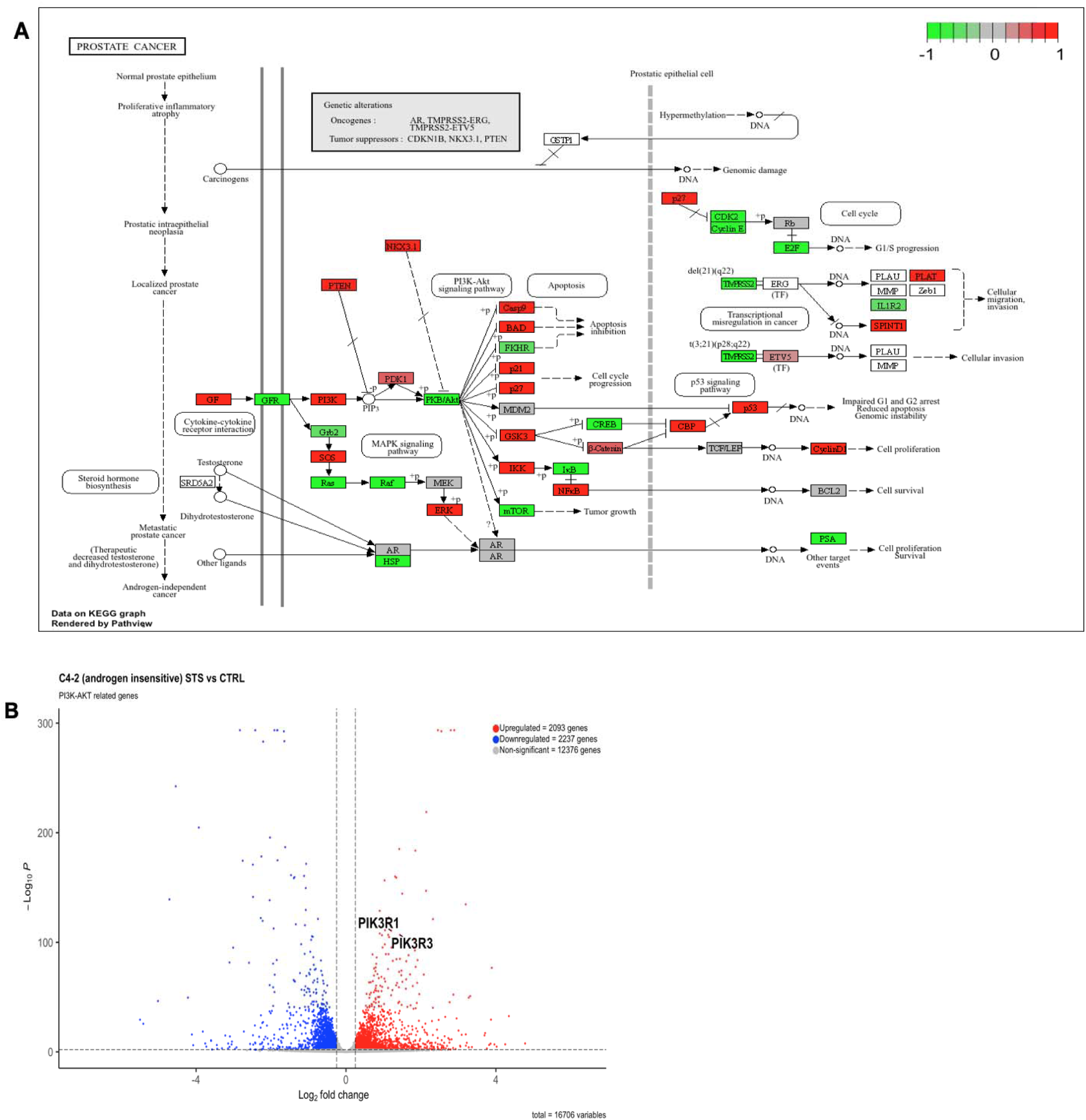
RNAseq and KEGG enrichment analysis to identify escape pathway mechanism. **A)** Significantly enriched prostate cancer KEGG pathways of C4-2 PCa grown in control or STS media. Red= KEGG node including up-regulated genes; green= KEGG node including down-regulated genes. STS vs. Ctrl. **B)** Volcano plot of the differential gene expression in androgen-independent C4-2 PCa grown in control or STS media. Highlighted are components of PI3K signaling pathways.

These studies provide a rationale explaining disappointing results of previous trials testing monotherapy Akt pathway inhibitors alone in advanced PCa^39^. The combination of PI3K inhibitors and androgen blockade can completely regress early-stage PCa in a PTEN loss mouse model^26,40^. However PI3K inhibitors have limited use due to adverse events incl. hyperglycemia. PI3K pathway inhibitors are more effective in decreasing PCa growth when combined with dietary modifications that decrease serum glucose and suppress insulin feedback^41^. Salvadori et al. has shown that FMD cycles reduce the hyperglycemic side effect of PI3K inhibitors^14^ while causing breast cancer tumor regression in combination with various kinase inhibitors^14,24^. We anticipate a synergistic effect with castration in repressing PCa tumor growth^13^. In fact, fasting/FMD might cooperate with hormone therapy and AKT inhibition by blocking AR or AKT signaling further, or blocking additional escape pathways such as insulin or leptin signaling. Notably in our work we showed that the combination of fasting mimicking medium conditions and simvastatin causes a major decrease in AR levels (**Fig.9**).

Initially, we established dose response curves for Pictilisib based on literature research to identify their effective dose. Several clinical trials have demonstrated the benefits of pictilisib, a pharmacological PI3K-AKT-mTOR inhibitor, in various types of cancer^42,43^. However, due to side effects such as hyperglycemia, the FDA did not approve this drug. Alpelisib is another pharmacological PI3K inhibitor that, due to better tolerability and fewer side effects, has been approved by the FDA for certain cancers, including breast cancer. Therefore, we decided to adopt alpelisib instead of pictilisib for our experiments. **Figure. 11** shows the dose response with alpelisib in both hormone-sensitive and hormone-insensitive prostate cancer cell lines.

**Figure 11.**
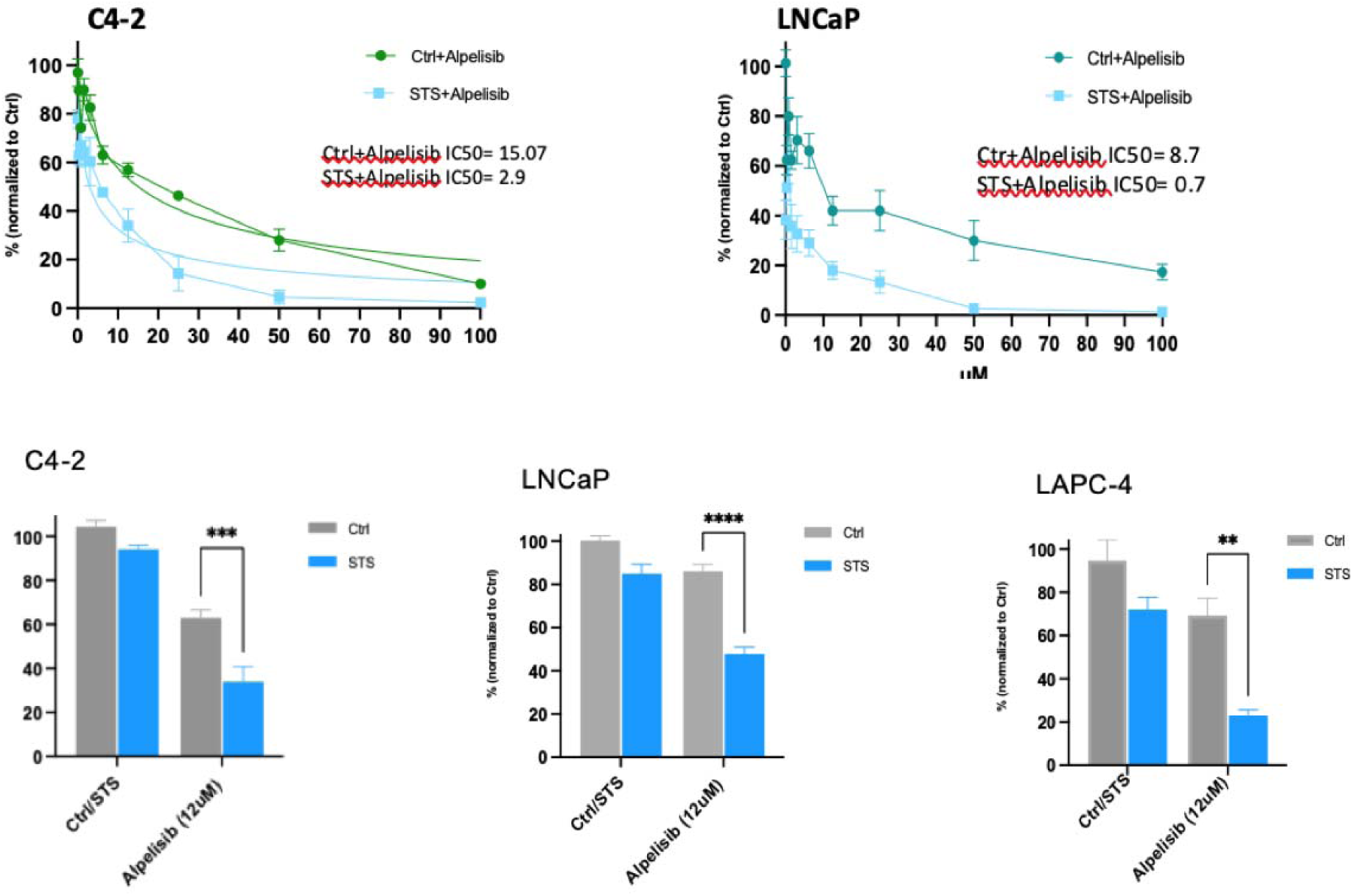
Combination targeting of PI3K signaling pathway with Alpelisib cooperate with fasting mimicking media/STS to kill prostate cancer cells. Prostate cancer cell lines were seeded in 96-well plates and were grown in CTRL (1g/l glucose,I0% FBS) or fasting STS (0.5 g/l glucose, I%FMS) media, after 24h cells were exposed to a gradient of drug concentrations for 48 hours, ranging from 0-100uM for Alpelisib. Cell viability of **A)** C4-2 and **B)** LNcaP Pca cells lines was assessed by MTS. C) C4-2 , D) LNCap and E) LAPC-4 cells were treated for 72h w/ or w/o fasting/STS and for the last 48h w/ or w/o 12uM alpelisib. Thereafter, cell viability was measured.

As anticipated, the pharmacological inhibition of the PI3K-AKT and mTOR pathways potentiates the effect of fasting/FMD: *in vitro*, combining fasting with pictilisib (PI3K pathway inhibitor), Enzalutamide, an anti-androgen drug, and Simvastatin which we identified through RNAseq results in enhanced cancer cell death and reduction of cancer cell survival. We showed that triple combination of drugs combined to fasting mimicking medium further increases the cytotoxicity in different prostate cancer cell lines as shown in Fig 12. Overall, we found that triple pictilisib+enzalutamide+simvastatin treatment combined with fasting mimicking medium, but not single or double treatment, strongly reduced the cell survival and cell proliferation. Cell survival was with MTS (**Fig.12**).

**Fig 12.**
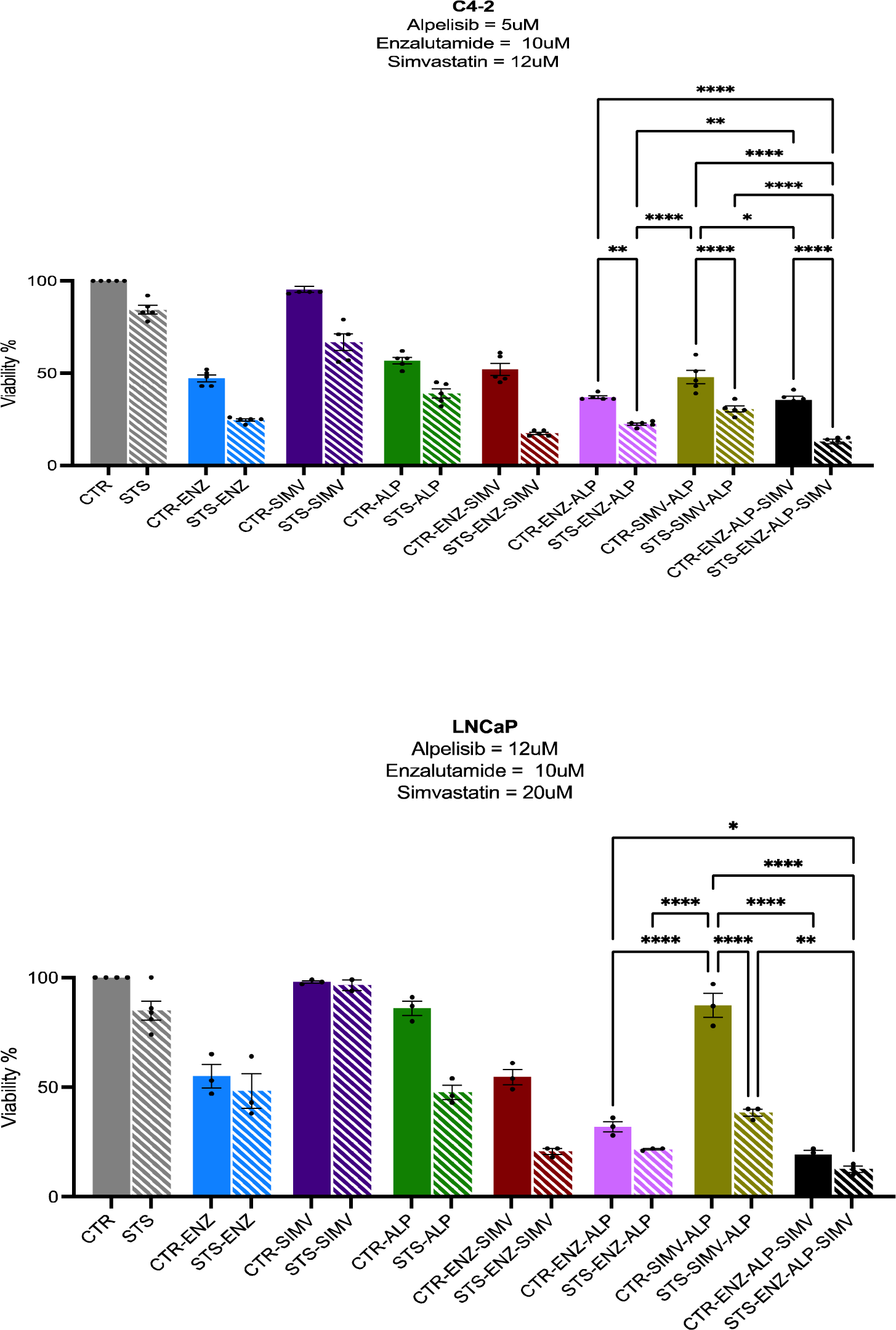
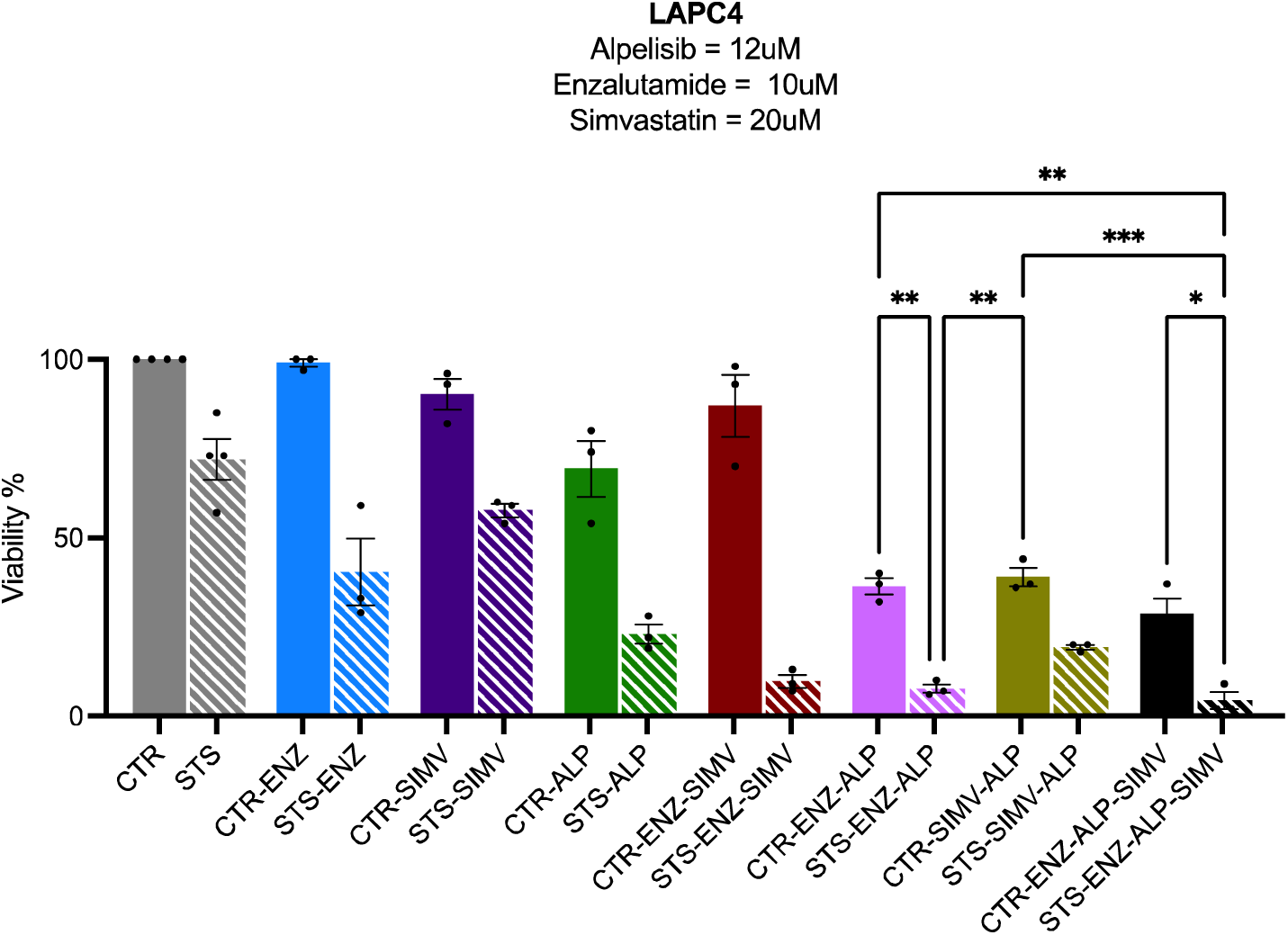
Fasting mimicking media/STS activates starvation escape mechanisms that can be targeted by drugs. **A)** C4-2, **B**) LNCap, and **C**) LAPC-4 were seeded in 96-well plates and cultured with or without STS media. After 24h cells were treated with Enzalutamide, alpelisib, and simvastatin at the indicated concentrations as single, dual, or triple treatment for 24hr and then cell viability was detected by MTS. Columns shows triplicate analysis. *,P<0.05 AND ***,P<0.001.

Based on our RNA-seq results, and on pervious data showing that by adding a cyclin-dependent kinase 4/6(CDK4/6) inhibitor such as palbociclib to fulvestrant combined with FMD not only prevents tumor growth for more than 160 days, but also leads to slow tumor shrinkage in a breast cancer xenograft model^13^, we evaluated the therapeutic effect of combining the CDK4/6 inhibitor palbociclib with alpelisib, simvastatin and enzalutamide. While palbociclib +alpelisib +simvastatin + enzalutamide significantly reduce cell viability, the addition of fasting mimicking medium to this pharmaoclogifcal combination further decrease the cancer survival and increase cell cytotoxicity in both hormone sensitive and hormone insensitive prostate cancer cells lines (**Fig 14**). In our previous experiment (**Fig. 7**), when simvastatin was used as a single agent in combination with FMM, a concentration of 30–40 μM was sufficient to reduce cell viability to nearly 0% in both androgen-sensitive and androgen-insensitive prostate cancer cell lines. However, when we tested drug combinations together with FMM, we lowered the concentration of simvastatin to 10–20 μM in order to better evaluate the effects of the other agents in combination with simvastatin. As a result, in **Fig. 8** and **Fig. 12**, cell viability decreased by about 70-80%, reflecting the lower simvastatin dose used in these combination experiments. All drug doses were adjusted and reduced in the combination experiments to allow evaluation of their potential synergistic effects.

**Figure 13.** Shows the upregulated cell cycle genes such as cyclin D1.

**Fig. 13.**
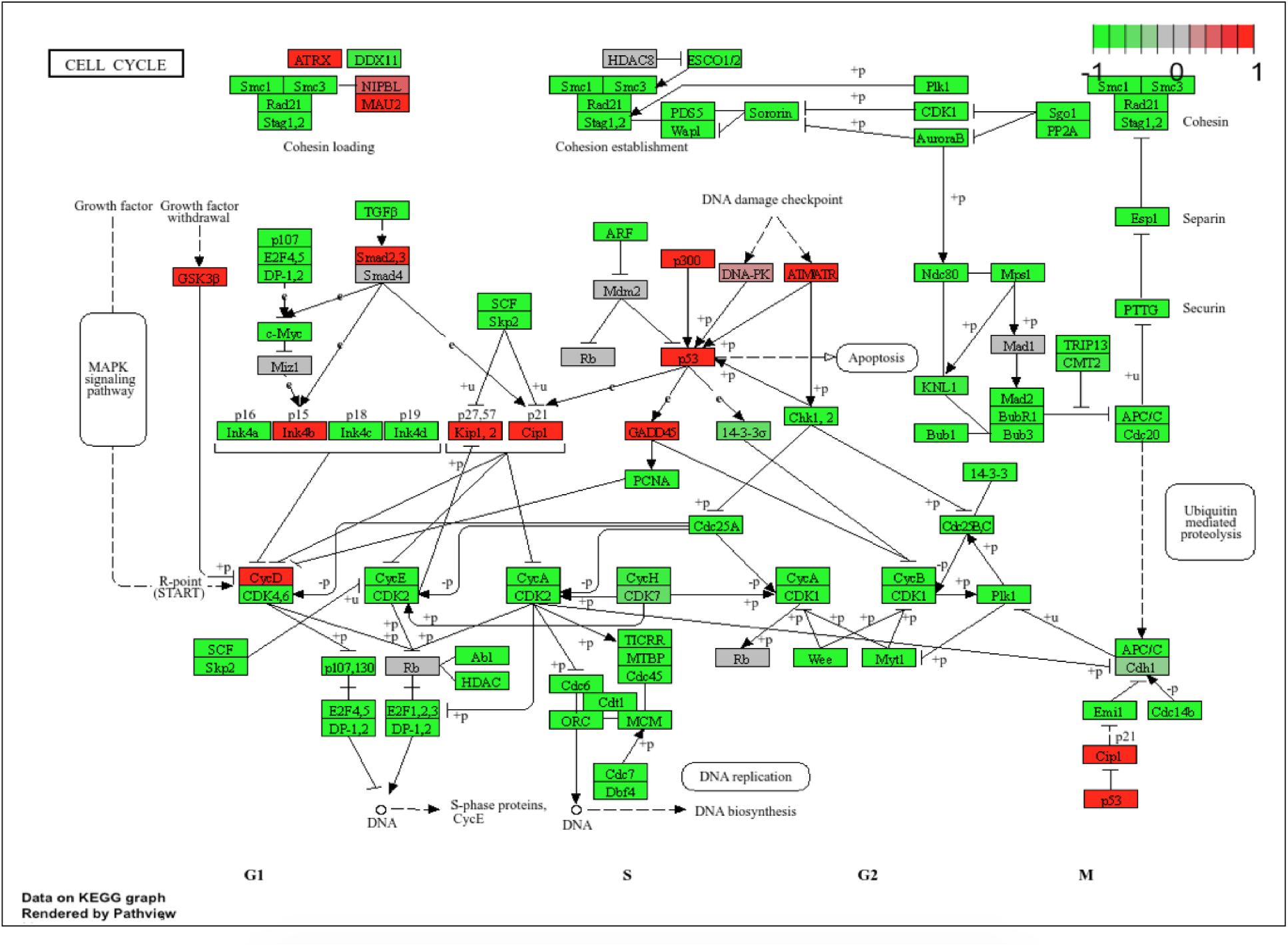
Targeting escape pathways to increase treatment efficacy. **A)** RNA sequencing on C4-2 cell line subjected to *in vitro* STS vs. CTR (for 48h) indicates up-regulation of cell cycle signaling via KEGG Pathview.

**Fig 14.**
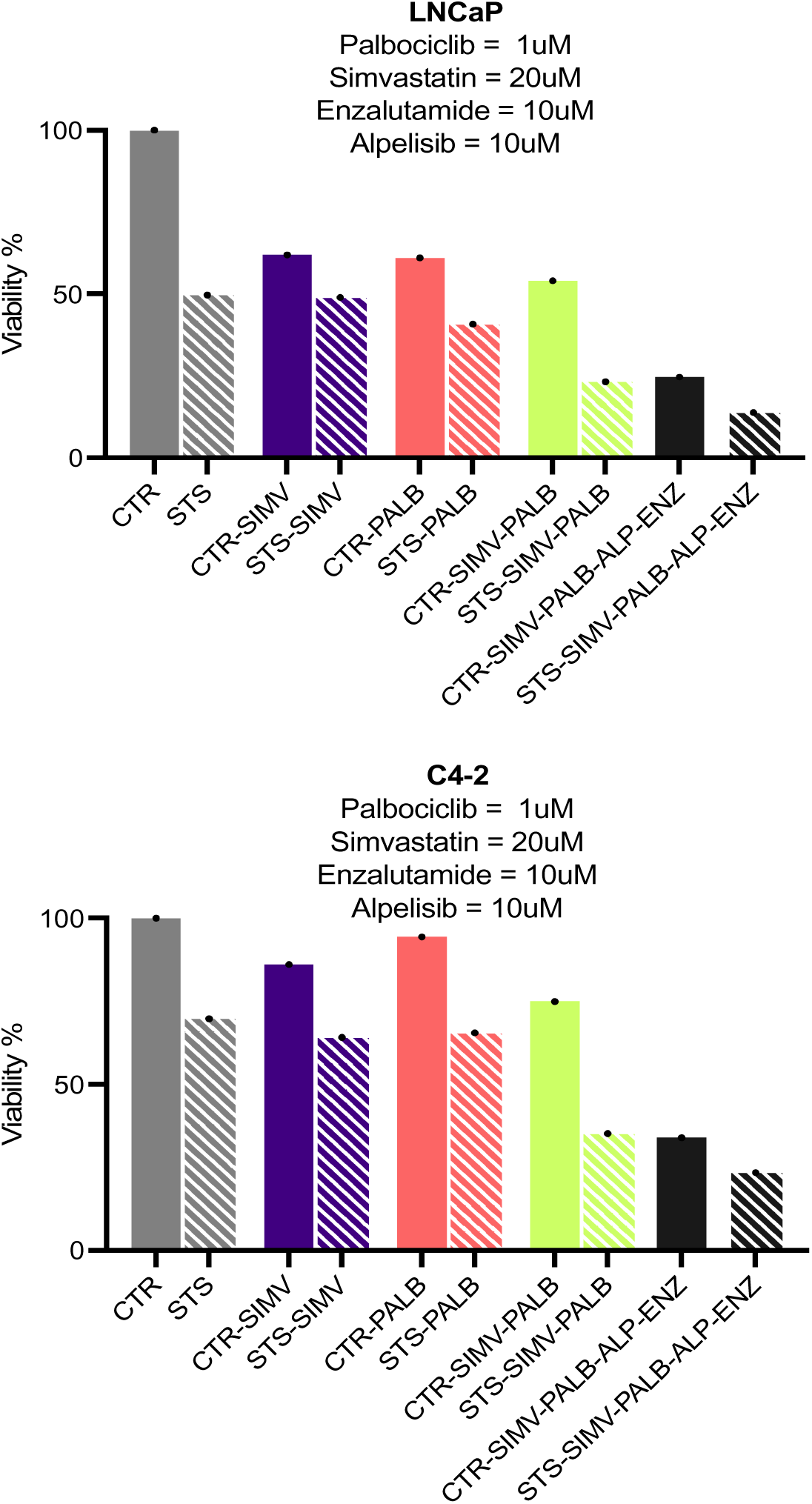
Targeting of escape pathways combined with anti-androgen potentiates fasting effects. LNCap and C4-2 cells were plated into 96-well plates and cultured with or without palbociclib (1uM), simvastatin(2OuM), enzalutamide (10uM), alpelisib (10uM) and STS conditions or their combinations for 48h. Cell viability was assessed by MTS. The result is representative of three independent experiments.

## Discussion

Our in vitro results indicate that fasting-mimicking conditions strongly sensitizes prostate cancer cells to multiple targeted agents. Under fasting mimicking conditions (FMM)/short-term starvation (STS), key oncogenic pathways such as PI3K–AKT and mTOR and cholesterol biosynthesis are either suppressed or activated. In fact, our RNA-seq data revealed upregulation of genes in cholesterol metabolism and putative “starvation escape” pathways, suggesting that nutrient stress triggers compensatory survival programs. This is in line with reports in other cancer models: for example, cyclic fasting increased expression of PI3K–AKT and mTOR and cell-cycle genes as a compensatory response in breast tumors, and inhibition of these pathways during FMD achieved synergistic tumor control^13,14^. In particular, the combination of fasting mimicking media (FMM) with PI3K, AKT, mTOR inhibitors led to markedly enhanced cell death in vitro and dramatically prolonged progression-free survival in xenograft models^14^. Similarly, agents targeting cholesterol biosynthesis (e.g. statins) become more cytotoxic under fasting in agreement with the activation of this pathways under nutrient scarce conditions. Fasting blunts circulating insulin, IGF-1 and leptin, downregulating the mevalonate pathway, which cooperates with statins to lower intracellular cholesterol, AKT/STAT3 signaling, and energy stores in tumors^31^. Thus, our observation that FMD augments the efficacy of simvastatin, rapamycin, PI3K inhibitors and enzalutamide likely reflects the requirement for PCa cells to turn to these pathways to survive under starvation conditions. Notably, simvastatin not only deprives cells of cholesterol precursors but also downregulates androgen receptor (AR) via mTOR inhibition^44^; in line with this, simvastatin plus enzalutamide dramatically suppressed enzalutamide-resistant cell growth in vitro and in vivo^44^. Moreover, dual blockade of AR and PI3K is known to produce synergistic anti-tumor effects in prostate cancer models^45^, which supports our finding of strong triple-combination synergy in vitro.

Our broader objective was to investigate whether combining in vitro data could help identify common escape pathways across multiple prostate cancer models, ultimately enabling the prediction of drug targets that synergize with the fasting-mimicking diet (FMD). Looking ahead, we aim to expand our understanding of how these escape pathways respond to FMD, standard-of-care therapies, or their combination by leveraging bioinformatics tools and in vivo models. Specifically, it will be valuable to utilize high-content bioinformatics approaches to integrate RNA-seq data with patient-derived genomic profiles, which may help identify individuals most likely to benefit from tailored FMD–drug regimens.

In summary, our work underscores the potential of exploiting differential responses of normal and PCa cells to FMD conditions which activate survival pathways which can be easily targeted with low toxicity drugs. In addition fasting/FMD conditions are well established to lower the toxicity of PI3K and mTOR-targeting drugs^14,46^ which could make it an ideal partner of standard of care prostate cancer drugs. A combination of refined models, integrated omics, and careful clinical strategies will be needed to fully realize the promise of FMD-based combination treatments.

## Methods

### Cell lines and cell culture

The human prostate cancer cell lines LNCaP and C4-2 was purchased from American Type Culture Collection (ATCC); the LAPC-4 was gifted from Dr. Henning at UCLA; 22RV-1 was gifted from Dr. Cohen at USC. Although LNCaP cells are androgen-dependent, C4-2 cells were derived from LNCaP but are androgen-independent. All the cells were maintained in RPMI 1640 (Corning) supplemented with 10% fetal bovine serum and 1% penicillin. Cultured cells were maintained in a humidified, 5% CO2 atmosphere at 37 °C and were passaged twice a week. In vitro, Short-Term starvation medium (STS) consists in a RPMI-1640 without glucose (RPMI-1640 no glucose, Life Technologies Cat# 11966025) supplemented with 0.5 g/l glucose (Sigma-Aldrich, Cat.#G8769) and 1% FBS. Control medium(CTR) consists in a RPMI-1640 no glucose medium supplemented with 1 g/l glucose and 10% FB.

### Cell Viability and Proliferation assay

Cell proliferation was determined at 24, 48, and 72 hours by colorimetric MTS assay (G3582, Promega). Briefly, cells will be seeded in 96-well plates at 20,000 cells/well, respectively, in complete media in triplicates to allow adhesion for 24 hours. On the following day, the medium will be replaced with either control (10% FBS, 1.0 g/L glucose) or STS (1% FBS, 0.5 g/L glucose) media. After 24 to 72 hours, cell proliferation was analyzed by adding 20 μl of MTS solution. After 4Lhr incubation at 37°C and 5% CO_2_, the absorbance of formazan product measured with a microplate reader at 490Lnm which is directly proportional to the cell count to quantify living cells in culture. Viability presented as a mean of MTS reduction level of treated cells from control cells.

Cell growth was assessed using crystal violet assay. Briefly, cells were plated in 96-well plates and allowed to attach for 48h before treatments. The cells were treated with tested drugs for 24 to 72 hours. Growth medium including the tested drugs were renewed every day. After treatments, the cells were stained with 50uL crystal violet for 20 min. After washing three times with water, 200ul methanol added to each well. Absorbance at 590 nm wavelength were measured with a microplate reader. Each sample was assayed in triplicate in three independent experiments.

### Protein extraction and Western Blot Analysis

For protein extraction, cells were washed in ice-cold PBS and lysates were prepared in radioimmunoprecipitation assay (RIPA) lysis buffer supplemented with protease and phosphatase inhibitors (#78446, Thermo Fisher). The lysates were shaken by vortex for 15 min and centrifuged at 15000 rpm for 15 min at 4 °C. After quantification, proteins (20-40µg) were separated by 4-20% SDS-PAGE, and transferred to a PVDF membrane (Immobilon-P, Millipore S.p.A). The following primary antibodies were used for protein detection: anti-AKT and anti-phospho-Ser 473 Akt (#9272s,#4058s), anti-p70 S6K and anti-phospho-Thr 389 p70 S6K (9202,#9206s), anti-CCND1 (#2978s), GAPDH (#2118L), (all from Cell Signaling Technology). Antibody against SREBP-2 (sc-5603) and AR (sc-7305) were obtained from Santa Cruz Biotechnology. After binding with primary antibody, the membrane was washed with TBS three times for 5 minutes, then incubated with anti-Mouse or anti-Rabbit secondary antibody for 1 hour at room temperature. Final detection was done with enhanced chemiluminescent reagents. Quantitative measurements of Western blot analysis were performed using GraphPad software (Prism 7).

### RNA Extraction

Total RNA will be isolated using the miRNeasy Mini Kit (#74124, QIAGEN) according to the manufacturer’s protocol.

### RNA sequencing

Libraries for RNAseq were prepared following the manufacturer protocols for transcriptome sequencing with the Illumina NextSeq 550DX sequencer (ILLUMINA). Total RNA were isolated from cells, using the miRNeasy Mini Kit (QIAGEN), according to the manufacturer’s instructions. Following sequencing, the raw data were aligned to mouse mm10 genome using Bowtie2 algorithm in Galaxy server. Gene counts will be then processed in R software for differential gene expression and downstream enrichment analysis. Briefly, the data were imported in R and genes without expression (total gene counts among the samples equal to zero) were excluded from the analysis. Then, the expression data underwent differential gene expression using DESeq2 R package^47^. The identified differentially expressed genes were analyzed using Ingenuity Pathway Analysis software and other pathway enrichment analysis platforms such as gProfiler, Gorilla, and Enrichr^48–52^. Heatmaps and Volcano plots were generated respectively with R packages ‘pheatmap’ and ‘EnhancedVolcano’, and gene set enrichment analysis (GSEA^53^). GO:BP visualization carried out in cytoscape using the EnrichmentMap and autoannotate plugins^54,55^

### Reagent’s preparation

#### Alpelisib

Alpelisib was purchased from MedchemTronica (Cat. #: HY-15244) and was dissolved in DMSO. Stock solutions of 10mM were prepared for *in vitro* experiments and stored at −80°C.

#### Rapamycin

Rapamycin was purchased from MedchemTronica (Cat. #: HY-10219) and was dissolved in DMSO. Stock solutions of 20mM and 20mg/mL were prepared for in vitro and in vivo experiments, respectively, and stored at −80°C.

#### Pictilisib

Pictilisb was purchased from MedchemTronica (Cat. #: HY-50094) and was dissolved in DMSO. Stock solutions of 10mM and 200mg/mL were prepared for in vitro and in vivo experiments, respectively, and stored at −80°C.

#### Enzalutamide

Enzalutamide was purchased from Selleckchem (Cat#: S1250) and was dissolved in DMSO. Stock solution of 10mM and 100 mg/ml were prepared for in vitro and in vivo experiments, respectively, and stored at −80°C.

#### Simvastatin

Simvastatin was purchased from APExBIO (Cat#: A8522) and was dissolved in DMSO. Stock solution of 10mM and 200 mg/ml were prepared for in vitro and in vivo experiments, respectively, and stored at −80°C.

### Statistics

Group differences means (± SEM) were analyzed by one-way ANOVA with Fisher’s LSD test, Kruskal-Wallis with Uncorrected Dunn’s test for nonparametric distributions, or Welch’s ANOVA when variances were unequal across groups. Significance was defined as pL<L0.05. Analyses used GraphPad Prism version 10 (GraphPad Software, San Diego, CA).

## Notes

### Competing Interest Statement

The authors have declared no competing interest.

## References

1 Sung, H. et al. Global Cancer Statistics 2020: GLOBOCAN Estimates of Incidence and Mortality Worldwide for 36 Cancers in 185 Countries. CA Cancer J Clin 71, 209–249, doi:10.3322/caac.21660 (2021).

2 Dai, C., Heemers, H. & Sharifi, N. Androgen Signaling in Prostate Cancer. Csh Perspect Med 7, a030452, doi:10.1101/cshperspect.a030452 PMID - 28389515 (2017).

3 Rebello, R. J. et al. Prostate cancer. Nat Rev Dis Primers 7, 9, doi:10.1038/s41572-020-0243-0 PMID - 33542230 (2021).

4 Kaarbo, M. et al. PI3K-AKT-mTOR pathway is dominant over androgen receptor signaling in prostate cancer cells. Cell Oncol 32, 11–27, doi:10.3233/CLO-2009-0487 (2010).

5 Squillace, R. M. et al. Synergistic activity of the mTOR inhibitor ridaforolimus and the antiandrogen bicalutamide in prostate cancer models. Int J Oncol 41, 425–432, doi:10.3892/ijo.2012.1487 (2012).

6 Manin, M. et al. Androgen receptor expression is regulated by the phosphoinositide 3-kinase/Akt pathway in normal and tumoral epithelial cells. Biochem J 366, 729–736, doi:10.1042/BJ20020585 (2002).

7 Noda, T. et al. Longterm exposure to leptin enhances the growth of prostate cancer cells. Int J Oncol 46, 1535–1542, doi:10.3892/ijo.2015.2845 (2015).

8 Raffaghello, L. et al. Starvation-dependent differential stress resistance protects normal but not cancer cells against high-dose chemotherapy. Proceedings of the National Academy of Sciences of the United States of America 105, 8215–8220, doi:0708100105 [pii] 10.1073/pnas.0708100105 (2008).

9 Longo, V. D. & Mattson, M. P. Fasting: Molecular Mechanisms and Clinical Applications. Cell metabolism 19, 181–192, doi:10.1016/j.cmet.2013.12.008 (2014).

10 Lee, C. et al. Reduced levels of IGF-I mediate differential protection of normal and cancer cells in response to fasting and improve chemotherapeutic index. Cancer research 70, 1564–1572, doi:0008-5472.CAN-09-3228 [pii] 10.1158/0008-5472.CAN-09-3228 (2010).

11 Safdie, F. M. et al. Fasting and cancer treatment in humans: A case series report. Aging 1, 988–1007 (2009).

12 Biase, S. D. et al. Fasting regulates EGR1 and protects from glucose-and dexamethasone-dependent sensitization to chemotherapy. Plos Biol 15, e2001951, doi:10.1371/journal.pbio.2001951 PMID - 28358805 (2017).

13 Caffa, I. et al. Fasting-mimicking diet and hormone therapy induce breast cancer regression. Nature 583, 620–624, doi:10.1038/s41586-020-2502-7 (2020).

14 Salvadori, G. et al. Fasting-mimicking diet blocks triple-negative breast cancer and cancer stem cell escape. Cell Metab 33, 2247–2259 e2246, doi:10.1016/j.cmet.2021.10.008 (2021).

15 de Bono, J. S. et al. Randomized Phase II Study Evaluating Akt Blockade with Ipatasertib, in Combination with Abiraterone, in Patients with Metastatic Prostate Cancer with and without PTEN Loss. Clin Cancer Res 25, 928–936, doi:10.1158/1078-0432.CCR-18-0981 (2019).

16 Nguyen, C., Lairson, D. R., Swartz, M. D. & Du, X. L. Risks of Major Long-Term Side Effects Associated with Androgen-Deprivation Therapy in Men with Prostate Cancer. Pharmacotherapy 38, 999–1009, doi:10.1002/phar.2168 (2018).

17 de Groot, S. et al. Fasting mimicking diet as an adjunct to neoadjuvant chemotherapy for breast cancer in the multicentre randomized phase 2 DIRECT trial. Nature communications 11, 3083, doi:10.1038/s41467-020-16138-3 (2020).

18 Di Biase, S. et al. Fasting-Mimicking Diet Reduces HO-1 to Promote T Cell-Mediated Tumor Cytotoxicity. Cancer cell 30, 136–146, doi:10.1016/j.ccell.2016.06.005 (2016).

19 Di Tano, M. et al. Synergistic effect of fasting-mimicking diet and vitamin C against KRAS mutated cancers. Nature communications 11, 2332, doi:10.1038/s41467-020-16243-3 (2020).

20 Di Biase, S. et al. Fasting-Mimicking Diet Reduces HO-1 to Promote T Cell-Mediated Tumor Cytotoxicity. Cancer Cell 30, 136–146, doi:10.1016/j.ccell.2016.06.005 PMID - 27411588 (2016).

21 Vernieri, C. et al. Exploiting FAsting-mimicking Diet and MEtformin to Improve the Efficacy of Platinum-pemetrexed Chemotherapy in Advanced LKB1-inactivated Lung Adenocarcinoma: The FAME Trial. Clinical lung cancer, doi:10.1016/j.cllc.2018.12.011 (2018).

22 Cortellino, S. & Longo, V. D. Metabolites and Immune Response in Tumor Microenvironments. Cancers (Basel) 15, doi:10.3390/cancers15153898 (2023).

23 Cortellino, S. et al. Fasting renders immunotherapy effective against low-immunogenic breast cancer while reducing side effects. Cell Rep 40, 111256, doi:10.1016/j.celrep.2022.111256 (2022).

24 Vernieri, C. et al. Fasting-Mimicking Diet Is Safe and Reshapes Metabolism and Antitumor Immunity in Patients with Cancer. Cancer Discov 12, 90–107, doi:10.1158/2159-8290.Cd-21-0030 (2022).

25 Wu, H. C. et al. Derivation of androgen-independent human LNCaP prostatic cancer cell sublines: role of bone stromal cells. Int J Cancer 57, 406–412, doi:10.1002/ijc.2910570319 (1994).

26 Mulholland, D. J. et al. Cell autonomous role of PTEN in regulating castration-resistant prostate cancer growth. Cancer Cell 19, 792–804, doi:10.1016/j.ccr.2011.05.006 (2011).

27 Wang, Y. et al. Regulation of androgen receptor transcriptional activity by rapamycin in prostate cancer cell proliferation and survival. Oncogene 27, 7106–7117, doi:10.1038/onc.2008.318 PMID - 18776922 (2008).

28 Gkioni, L. et al. The geroprotectors trametinib and rapamycin combine additively to extend mouse healthspan and lifespan. Nat Aging 5, 1249–1265, doi:10.1038/s43587-025-00876-4 (2025).

29 Selvarani, R., Mohammed, S. & Richardson, A. Effect of rapamycin on aging and age-related diseases-past and future. Geroscience 43, 1135–1158, doi:10.1007/s11357-020-00274-1 (2021).

30 Pushpakom, S. et al. Drug repurposing: progress, challenges and recommendations. Nat Rev Drug Discov 18, 41–58, doi:10.1038/nrd.2018.168 (2019).

31 Khalifa, A. et al. Cyclic fasting bolsters cholesterol biosynthesis inhibitors’ anticancer activity. Nat Commun 14, 6951, doi:10.1038/s41467-023-42652-1 (2023).

32 Iannelli, F. et al. Targeting Mevalonate Pathway in Cancer Treatment: Repurposing of Statins. Recent Pat Anticancer Drug Discov 13, 184–200, doi:10.2174/1574892812666171129141211 (2018).

33 Zhai, C. et al. Efficacy of statin treatment based on cardiovascular outcomes in elderly patients: a standard meta-analysis and Bayesian network analysis. J Int Med Res 48, 300060520926349, doi:10.1177/0300060520926349 (2020).

34 Gordon, J. A. et al. Statin use and survival in patients with metastatic castration-resistant prostate cancer treated with abiraterone or enzalutamide after docetaxel failure: the international retrospective observational STABEN study. Oncotarget 9, 19861–19873, doi:10.18632/oncotarget.24888 (2018).

35 Kochuparambil, S. T., Al-Husein, B., Goc, A., Soliman, S. & Somanath, P. R. Anticancer efficacy of simvastatin on prostate cancer cells and tumor xenografts is associated with inhibition of Akt and reduced prostate-specific antigen expression. J Pharmacol Exp Ther 336, 496–505, doi:10.1124/jpet.110.174870 (2011).

36 Cham, J., Venkateswaran, A. R. & Bhangoo, M. Targeting the PI3K-AKT-mTOR Pathway in Castration Resistant Prostate Cancer: A Review Article. Clinical genitourinary cancer 19, 563 e561–563 e567, doi:10.1016/j.clgc.2021.07.014 (2021).

37 Toren, P. et al. Combination AZD5363 with Enzalutamide Significantly Delays Enzalutamide-resistant Prostate Cancer in Preclinical Models. Eur Urol 67, 986–990, doi:10.1016/j.eururo.2014.08.006 (2015).

38 Kolinsky, M. P. et al. A phase I dose-escalation study of enzalutamide in combination with the AKT inhibitor AZD5363 (capivasertib) in patients with metastatic castration-resistant prostate cancer. Ann Oncol 31, 619–625, doi:10.1016/j.annonc.2020.01.074 (2020).

39 Chee, K. G. et al. The AKT inhibitor perifosine in biochemically recurrent prostate cancer: a phase II California/Pittsburgh cancer consortium trial. Clin Genitourin Cancer 5, 433–437, doi:10.3816/CGC.2007.n.031 (2007).

40 Carver, Brett S. et al. Reciprocal Feedback Regulation of PI3K and Androgen Receptor Signaling in PTEN-Deficient Prostate Cancer. Cancer Cell 19, 575–586, doi:10.1016/j.ccr.2011.04.008 PMID - 21575859 (2011).

41 Hopkins, B. D. et al. Suppression of insulin feedback enhances the efficacy of PI3K inhibitors. Nature 560, 499–503, doi:10.1038/s41586-018-0343-4 PMID – 30051890 (2018).

42 Schmid, P. et al. Phase II Randomized Preoperative Window-of-Opportunity Study of the PI3K Inhibitor Pictilisib Plus Anastrozole Compared With Anastrozole Alone in Patients With Estrogen Receptor-Positive Breast Cancer. J Clin Oncol 34, 1987–1994, doi:10.1200/jco.2015.63.9179 (2016).

43 Sarker, D. et al. First-in-human phase I study of pictilisib (GDC-0941), a potent pan-class I phosphatidylinositol-3-kinase (PI3K) inhibitor, in patients with advanced solid tumors. Clin Cancer Res 21, 77–86, doi:10.1158/1078-0432.Ccr-14-0947 (2015).

44 Kong, Y. et al. Inhibition of cholesterol biosynthesis overcomes enzalutamide resistance in castration-resistant prostate cancer (CRPC). J Biol Chem 293, 14328–14341, doi:10.1074/jbc.RA118.004442 (2018).

45 Qi, W. et al. Reciprocal feedback inhibition of the androgen receptor and PI3K as a novel therapy for castrate-sensitive and -resistant prostate cancer. Oncotarget 6, 41976–41987, doi:10.18632/oncotarget.5659 PMID - 26506516 (2015).

46 Caffa, I. et al. Author Correction: Fasting-mimicking diet and hormone therapy induce breast cancer regression. Nature 588, E33, doi:10.1038/s41586-020-2957-6 (2020).

47 Love, M. I., Huber, W. & Anders, S. Moderated estimation of fold change and dispersion for RNA-seq data with DESeq2. Genome Biol 15, 550, doi:10.1186/s13059-014-0550-8 (2014).

48 Kolberg, L. et al. g:Profiler-interoperable web service for functional enrichment analysis and gene identifier mapping (2023 update). Nucleic Acids Res 51, W207–w212, doi:10.1093/nar/gkad347 (2023).

49 Eden, E., Navon, R., Steinfeld, I., Lipson, D. & Yakhini, Z. GOrilla: a tool for discovery and visualization of enriched GO terms in ranked gene lists. BMC Bioinformatics 10, 48, doi:10.1186/1471-2105-10-48 (2009).

50 Kuleshov, M. V. et al. Enrichr: a comprehensive gene set enrichment analysis web server 2016 update. Nucleic Acids Res 44, W90–97, doi:10.1093/nar/gkw377 (2016).

51 Xie, Z. et al. Gene Set Knowledge Discovery with Enrichr. Curr Protoc 1, e90, doi:10.1002/cpz1.90 (2021).

52 Chen, E. Y. et al. Enrichr: interactive and collaborative HTML5 gene list enrichment analysis tool. BMC Bioinformatics 14, 128, doi:10.1186/1471-2105-14-128 (2013).

53 Subramanian, A. et al. Gene set enrichment analysis: a knowledge-based approach for interpreting genome-wide expression profiles. Proc Natl Acad Sci U S A 102, 15545–15550, doi:10.1073/pnas.0506580102 (2005).

54 Shannon, P. et al. Cytoscape: a software environment for integrated models of biomolecular interaction networks. Genome Res 13, 2498–2504, doi:10.1101/gr.1239303 (2003).

55 Merico, D., Isserlin, R., Stueker, O., Emili, A. & Bader, G. D. Enrichment map: a network-based method for gene-set enrichment visualization and interpretation. PLoS One 5, e13984, doi:10.1371/journal.pone.0013984 (2010).

